# Neural signatures of competition between voluntary and involuntary influences over the focus of attention in visual working memory

**DOI:** 10.1101/2023.10.05.561134

**Authors:** Yun Ding, Bradley R. Postle, Freek van Ede

## Abstract

Adaptive behaviour relies on the selection and prioritisation of relevant sensory inputs from the external environment as well as from among internal sensory representations held in working memory. Recent behavioural evidence suggests that the classic distinction between voluntary (goal-driven) and involuntary (stimulus-driven) influences over attentional allocation also applies to the selection of internal representations held in working memory. In the current EEG study, we set out to investigate the neural dynamics associated with the competition between voluntary and involuntary control over the focus of attention in visual working memory. We show that when voluntary and involuntary factors compete for the internal focus of attention, prioritisation of the appropriate item is delayed – as reflected both in delayed gaze biases that track internal selection and in delayed neural beta (15-25 Hz) dynamics that track the planning for the upcoming memory-guided manual action. We further show how this competition is paralleled – possibly resolved – by an increase in frontal midline theta (4-8 Hz) activity that, moreover, predicts the speed of ensuing memory-guided behaviour. Finally, because theta increased following retrocues that effectively reduced working-memory load, our data unveil how frontal theta activity during internal attentional focusing tracks demands on cognitive control over and above working-memory load. Together, these data yield new insight into the neural dynamics that govern the focus of attention in visual working memory, and disentangles the contributions of frontal midline theta activity to the processes of control versus retention in working memory.

## Introduction

Working memory refers to the ability to maintain and manipulate relevant information for the guidance of perception, thought, and action (Baddeley, 1992; D’Esposito & Postle, 2015). Previous studies suggest that working-memory and attention interact to guide ensuing memory-guided behavior (Olivers et al., 2006; Theeuwes et al., 2009; Woodman et al., 2013; Ding et al., 2019; for reviews, see Olivers et al., 2011; van Ede & Nobre, 2023). For example, one influential line of work has demonstrated that information held in working memory can capture attention in a manner similar to the involuntary capture of attention by distracting visual stimuli (Soto et al., 2005; Olivers et al., 2006; Bahle et al., 2018; For a review, see Soto et al., 2008). A complementary line of research has focused on the role of attention in selecting and prioritising the contents of mind themselves (Griffin & Nobre, 2003; see Souza & Oberauer, 2016; van Ede & Nobre, 2023 for reviews). In a recent study, van Ede et al. (2020) used eye-tracking to demonstrate that the selection and prioritisation of one from among multiple items in visual working memory can be driven not only by voluntary but also involuntary factors (paralleling the classic distinction between voluntary and involuntary influences over externally directed selective attention, c.f., Corbetta & Shulman, 2002; Dalvit & Eimer, 2011; Yantis & Jonides, 1990; Posner, 2016). In the present study, we replicated the procedures of van Ede et al. (2020) while additionally recording the electroencephalogram (EEG). This enabled us to assess the dynamic neural bases of voluntary and involuntary influences over the focus of attention inside visual working memory – as well as of the neural processes involved in ensuring appropriate attentional allocation when voluntary and involuntary influence compete for the internal focus of attention.

In the study that motivated this one (van Ede et al., 2020), participants encoded the orientation of two bars, one presented on each side of fixation, and each in a different color. During the working-memory delay, the central fixation cross flashed briefly in one of the two colors serving as a retrocue (Griffin & Nobre, 2003; Souza & Oberauer, 2016; van Ede & Nobre, 2023). During “pro-cue” blocks, this retrocue color indicated (with 100% validity) the item that would be tested, whereas during “anti-cue” blocks it indicated (with 100% validity) that the other item would be tested. In “null-cue” blocks, as a control condition, the retrocue did not predict which item would be tested (50% validity). Thus, both the pro- and anti-retrocues were informative, allowing for internal selection of one of the two items, and a concomitant decrease of working-memory load. Null-retrocues, in contrast, were non-informative and did not change working-memory load. Results from critical anti-cue blocks showed a delayed gaze bias – as an index of internal attentional allocation (van Ede et al., 2019; Liu et al., 2022) – toward the cued (but color-incongruent) item, suggesting that an initial involuntary shift of attention (toward the color-incongruent item) was overcome by a later-developing voluntary deployment to the contextually appropriate memory item.

In the present experiment, we took this task to EEG to address several open questions regarding the neural dynamics supporting selective attentional prioritisation of working-memory content in the presence of both voluntary and involuntary factors. As a sanity check, we first inspected early visually event-related potentials (ERPs) time-locked to the retrocue to confirm that informative retrocues are processed differently than uninformative retrocues. Next, we investigated spectral dynamics in central motor electrodes to assess the time course of motor preparatory activity that has been reported to occur alongside visual retention in working-memory tasks (Nasrawi et al., 2023; Boettcher et al., 2021; Nasrawi & van Ede, 2022; Schneider et al., 2017; van Ede, Chekroud, Stokes, et al., 2019; Rösner et al., 2022), and that some hold to be a fundamental contributor to visual working memory (e.g., Postle, 2006; Postle et al., 2006; Postle & Hamidi, 2007). Such motor preparation signals are expected to follow, and hence index, the time course of item prioritization within working memory. Like what was previously observed for gaze, we hypothesised that such motor preparation signals would also be delayed following anti-compared to pro-retrocues, consistent with first needing to overcome the competition between involuntary and voluntary factors following anti-, but not pro-retrocues. Finally, of particular interest was oscillatory power in the theta-band (approximately 4-8 Hz) at frontal midline electrodes (“frontal-midline theta”, hereafter FMT), which has been linked to two aspects of working memory that we could isolate in this study: cognitive control and working memory load.

With regard to control, FMT is known to covary with demands on control (e.g., Cavanagh & Frank, 2014; Hsieh & Ranganath, 2014), such as with the requirement to reorganize the contents of working memory into a canonical order (Griesmayr et al., 2010). Raghavachari and colleagues (2001) observed that the power of FMT increases at the beginning of a working memory trial, is sustained at an elevated level through the retention period, then falls off precipitously at the end of the trial, a pattern interpreted as evidence for a role in “cognitive gating”. Additionally, however, FMT has also been seen to covary with working-memory load, consistent with a role in representation/retention per se. For example, Gevins et al., (1997) observed that FMT power was parametrically modulated by the number of items held in working memory (replicated by, e.g.: Jensen and Tesche, 2002; Zakrzewska & Brzezicka, 2014; Meltzer et al., 2008; van Ede et al., 2017). Consistently, brain stimulation (via transcranial magnetic stimulation or transcranial alternating-current stimulation) delivered in theta range to frontal midline cortex has been shown to improve working-memory performance (Riddle et al., 2020; Zakrzewska & Brzezicka, 2014; Reinhart and Nguyen, 2019). Although these two functions – control and retention – need not be mutually exclusive, in our design they can be dissociated: whereas anti-retrocue trials require control of the conflict between involuntarily and voluntarily triggered attention (relative to pro-retrocue trials), null-retrocue trials require the retention of more items (relative to pro- and anti-retrocue trials). Thus, our experiment enabled us to disentangle the involvement of FMT in these theoretically distinct functions.

## Methods

The experimental procedures were reviewed and approved by the ethical committee of the Faculty of Behavioral and Movement sciences Vrije Universiteit Amsterdam.

### Participants

A total of 27 participants (ranged 20-30 years; all right-handed) were recruited to achieve the predetermined sample size of 25, which was selected to match the sample-size in van Ede et al. (2020). One participant was replaced because the participant did not complete the experimental session, and a second due to excessive movement-related artifacts in the EEG data. All participants provided written informed consent before participation and reported normal or corrected-to-normal vision and no history of neurological disorder.

### Setup

The experiment was run on a PC (with a 1920x1080 pixel LCD monitor with a 240 Hz refresh rate) in a dimly-lit room, and the task programmed in Presentation. A chin rest was used to maintain a viewing distance of approximately 75 cm. Gaze position was tracked and recorded at a sampling rate of 1000 Hz with an Eyelink 1000 eye tracker (SR Research Ltd. Ottawa ON). The eye tracker was calibrated and validated for each participant before the experiment using a built-in 9-point calibration routine. Manual responses were made via computer mouse.

EEG signals were recorded with a 64-channel Biosemi system (1024 Hz sampling rate), with active electrodes distributed across the scalp using the international 10-20 positioning system. The left mastoid was used as the online reference and the right mastoid was used to derive an average mastoid reference offline. Two electro-oculography (EOG) electrodes were placed below and to the right of the right eye to monitor for eye movements and blinks.

### Task and procedure

The current experiment directly built on the task and procedure from van Ede et al. (2020) and thus had nearly identical procedures. In comparison to the original study, we made three adaptations. First and foremost, we here included neural EEG measurements. Second, to help disambiguate the retrocue from the memory probe, we adapted the retrocue to take the form of a colour patch presented behind the fixation cross rather than a change in the colour of the fixation cross itself (as we had used in the original study). Finally, to boost trial numbers in the most relevant conditions, we no longer included trials with grey neutral retrocues that we had included in the original study.

Each trial began with the brief simultaneous presentation of two visual stimuli of different colour and orientation. The stimuli were rectangular bars (approximately 5.7 ° visual angle in width and 0.8° in height), and appeared centered at a distance of approximately 5.7° visual angle to the left or right of fixation. On each trial one bar was colored purple and one green (location randomized across trials) and each appeared at an orientation that could vary between 1° and 180°. 750 msec after sample offset, a retrocue (a disc colored purple or green, with a radius of approximately 0.2°) appeared behind the fixation cross for 250 ms. 1750 msec after retrocue offset, the fixation cross changed color to purple or green, prompting recall of the sample matching that color. Participants were instructed to maintain central fixation and hold the computer mouse stationary until they were ready to initiate their orientation recall response. Movement of the mouse triggered the onset of the response dial, which appeared displaying a randomly selected orientation, and participants adjusted the orientation of the dial to indicate their recall of the orientation of the probed sample, confirming their response by clicking the left button on the mouse. The response dial was a circle centered on the fixation symbol (radius of 5.7°) with two smaller circles (like handles on a steering wheel) centered on its circumference and separated by 180°. Participants were trained that the imaginary diameter connecting the two “handles” corresponded to the recalled orientation. Participants had an unlimited amount of time to initiate their response, then 2.5 sec to complete it. Upon mouse click (or timeout) participants received feedback about the accuracy of their response (200 ms; a digit ranging from 1 to 100 with “1” reflecting the maximum possible error of 90° and “100” reflecting a perfect report), followed by an intertrial interval that varied randomly between 500-800 msec.

The central manipulation was that we included three types of retrocue trials that were blocked by category: “pro-retrocue”, “anti-retrocue”, and “null-retrocue”. On pro-, anti-, and null-retrocue trials, the color of the retrocue predicted that the to-be-recalled memory item would be the colour-matching memory item with 100%, 0%, and 50% validity, respectively (i.e., in null blocks, the cue predicted with 100% validity that the other, non-colour-matching, memory item would be probed). The experiment comprised 16 pro-, 16 anti-, and 16 null-retrocue blocks, administered in clusters of three (one block of each condition, randomly determined order), with 16 trials per block. During the entirety of each block, the word “pro”, “anti”, or “null” was displayed at the top of the screen, so that participants could always remind themselves of how to use the cue. Participants initiated each block by pressing the enter key on the keyboard. Between each cluster of blocks the eye tracker was recalibrated. Prior to data collection, each participant practiced one pro-block, followed by one anti-block, then one null-block.

### Analysis of behavioral (manual response) data

Behavioral data were analyzed with two measures: response times and reproduction errors. Response times were defined as the duration from probe onset to response initiation (i.e., initial movement of the cursor). Reproduction errors were defined as the absolute angular difference between the probed memory orientation and the reproduced one. Then we used a one-way repeated-measures ANOVA with factors retrocue informativeness (i.e., pro, anti, and null) and follow-up paired-samples t-tests with these two dependent variables.

### Analysis of eye-tracking data

Eye-tracking data were converted from their original .edf format to .asc format using the EDF2ASC application which is bundled with Eyelink. Subsquent processing was carried out with custom code in MATLAB. The data were cleaned of blink-related artifacts by removing signal ±120 ms around each blink. As in previous studies (van Ede et al., 2019; 2020), biases in gaze following informative pro and anti-retrocues were quantified using a measure of “towardness”: capturing the bias in horizontal gaze position as a function of the cued items location at encoding. For both pro- and anti-retrocue trials, positive values were assigned to bias in the direction of the to-be attended memory item (i.e., for anti-cue trials this was always the item whose color did *not* match the cue). After towardness was calculated per time point, for visualization, trial-average gaze-position time courses were smoothed by a moving averaging kernel with 5 samples.

### Analysis of EEG data

#### Preprocessing

EEG data were processed and analyzed in MATLAB with FieldTrip (Oostenveld et al., 2011). During preprocessing, the data were epoched from 500 ms before memory array onset to 1000 ms after probe onset (using ft_definetrial). The epoched data were re-referenced to the average of both mastoids (using ft_preprocessing). Bad channels were interpolated with the two lateral neighboring electrodes. Next, eye-related artifacts were detected using independent component analysis (ICA; using ft_componentanalysis) with the FastICA algorithm (Hyvarinen, 1999). ICA components corresponding to artifacts, identified via correlation with the EOG signal and assessment of their topography, were removed from the data. To exclude all trials in which blinks may have interfered with processing retrocues, we removed trials in which the vertical EOG contained samples with amplitude higher than 2000 uV. Finally, we visually detected and discarded trials with obviously high variance by utilizing “ft_rejectvisual” function with the “summary” method. To increase topographical specificity, we conducted a surface Laplacian transform (using ft_scalpcurrentdensity) on the preprocessed data.

#### Electrode and Frequency-Band Selection

Channel and frequency-band selections for all presented time-frequency analyses were predetermined as follows. To explore manual response preparation, we focused on dynamics in the beta-band (15-25 Hz) at electrode C3, which was contralateral to the response hand (Neuper, Wörtz, & Pfurtscheller, 2006; Baker, 2007; Nasrawi & van Ede, 2022). To explore control-related demands of our task, power of frontal midline theta-band oscillations (Sauseng et al., 2010; Jensen & Tesche, 2002) was measured from 4-8 Hz at electrode AFz. Statistical analyses were done across the full time-frequency axes at the selected electrodes. In addition, we visualised topographies for which responses were averaged for the above predetermined frequency bands.

#### Time-Frequency Analysis

Time-frequency responses from 2 to 40 Hz were obtained in steps of 1 Hz using a short-time Fourier transform. Data were Hanning-tapered with a sliding time window of 300 ms, progressing in steps of 10 ms. To compare time-frequency responses of different conditions (say, conditions “a” and “b”), we normalized the difference as percentage changes: ((a - b)/(a + b)) × 100. We focused on the period around retrocue onset (-500 to 2000 ms) to study retrocue-induced neural modulations.

### Statistical Evaluation

Statistical evaluation of all the gaze-position and spectral EEG data was performed with cluster-based permutation testing (Maris & Oostenveld, 2007). These analyses were conducted on the time-courses by considering clusters in time and on the time-frequency responses by considering clusters in both time and frequency, using 1000 permutations and an alpha level of 0.025.

## Results

### Behavior: informative cues enhance performance

Patterns in response-onset times (RTs) and reproduction errors (Fig. 2A) confirmed that participants used informative retrocues to internally select and prioritise the appropriate memory item following both pro- and anti-retrocues. As anticipated, responses were initiated most quickly on pro-trials (M = 343 ms), followed by anti- (M = 417 ms) and null-retrocue trials (M = 642 ms; F(2, 48) = 68.66, P < 0.001; pro vs. null, t(24) = -11.94, P < 0.001, d = 2.58; anti vs. null, t(24) = -7.28, P < 0.001, d = 1.70; pro vs. anti t(24) = -3.19, P = 0.004, d = 0.64; Fig. 2A, Left). Consistent with the RT data, reproduction errors were smallest for pro-retrocue trials (M = 9.26°), followed by anti- (M = 9.96°) and null-retrocue trials (M = 10.40°; F(2, 48) = 5.92, P = 0.005; pro vs. null, t(24) = -3.36, P = 0.003, d = 0.64; anti vs. null, t(24) = -1.19, P = 0.25, d = 0.18; pro vs. anti, t(24) = -2.44, P = 0.02, d = 0.38. Fig. 2A, Right).

**Figure 1.**
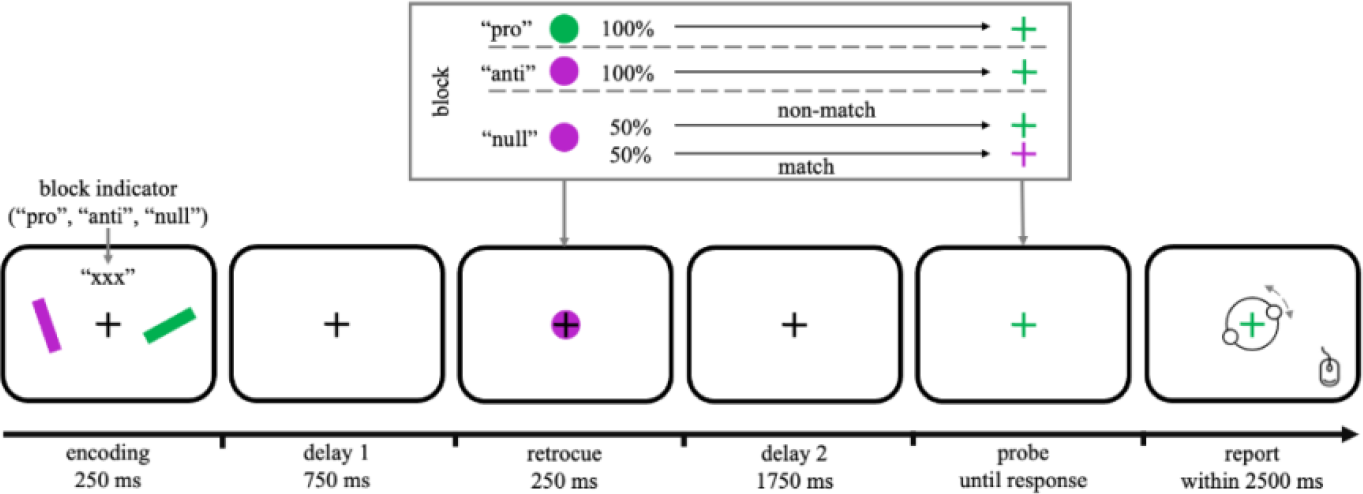
Visual working memory task with informative (pro, anti) and uninformative (null) retrocues. Participants encoded two visual tilted bars to reproduce the orientation of the probed memory item at the trial end. A dot colored with one of the memory colors was presented transiently at the fixation after the delay following the encoding, serving as the retrocue. After another delay, the central fixation cross was colored by one of the memory colors, serving as the probe for participants to report the matching item’s orientation. In the informative retrocue blocks, pro- and anti-retrocues were each 100% predictive of the to-be probed memory item but differed in whether their color also matched the probed item (pro) or matched the other item (anti). Null-retrocues were uninformative by matching the probed color or the other memory color for 50% of trials, respectively.

**Figure 2.**
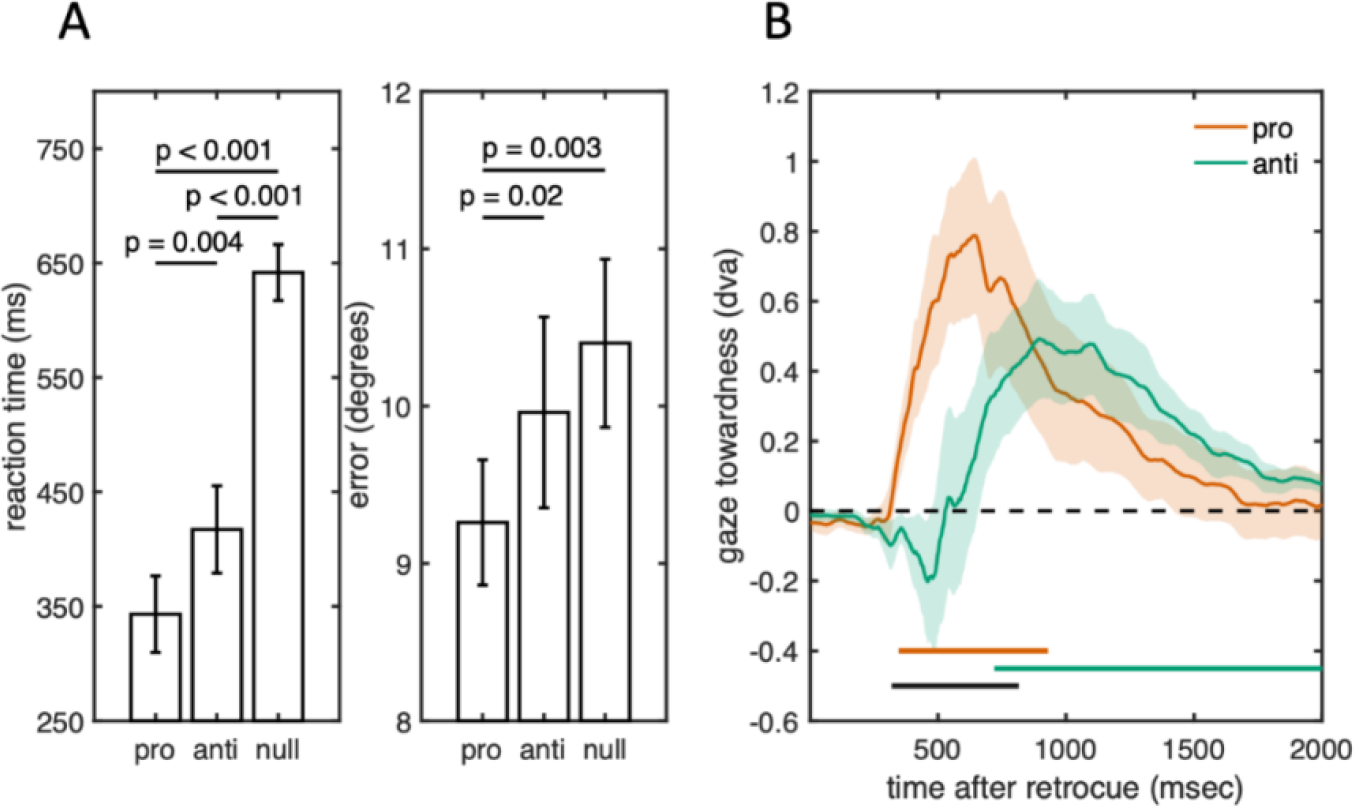
Behavioural performance and gaze biases of internal selective attention following pro- and anti-retrocues. (A) behavioral performance as a function of retrocues for internal selection. Error bars represent ±1 standard error. (B) pro- and anti-retrocues bias horizontal gaze toward memorized item locations. The red, green, and black horizontal lines indicate significant clusters for pro, anti, and the difference between them, respectively. The light shading around the time-courses in (B) indicate ±1 standard error.

### Eye tracking: gaze-bias signatures of selection and competition

Spatial biases in gaze following the retrocue (Fig. 2B) revealed signatures of selection in visual working memory following informative retrocues, in both the pro- and anti-retrocue conditions. In addition, gaze data confirmed competition between voluntary and involuntary factors revealed by the difference in selection dynamics following pro- and anti-retrocues.

In both conditions with informative retrocues, deviations of gaze from fixation could be observed beginning approximately 350 ms after retrocue onset, although the pattern differed markedly between the two. On pro-retrocue trials, beginning at around 350 ms, gaze bias shifted toward the location that had been occupied by the cued memory item, quickly differing from anti-retrocue trials (cluster P = 0.007) and (with a slightly longer latency) from baseline (cluster P = 0.012). This deviation reached its peak at approximately 650 ms after retrocue onset before reversing and becoming statistically indistinguishable from pre-cue fixation at approximately 1000 ms after retrocue onset. On anti-retrocue trials, in contrast, the early deviation of gaze was noisier and initially took on numerically (although not significantly) negative values before subsequently shifting toward the location of the appropriate item. That is, on anti-cue trials gaze bias initially trended toward the location of the sample whose color matched the retrocue, before subsequently shifting toward the memorised location of the relevant memory item. On anti-retrocue trials the (positively valued) gaze bias cluster was observed approximately 700 ms after cue onset (cluster P < 0.001), flattened out approximately 300 ms later, but extended for the remainder of the delay period.

The observed delay in gaze bias to the appropriate memory item following anti-compared to pro-retrocues is consistent with a competition between involuntary (colour-driven) and voluntary (goal-driven) factors over the focus of internal selective attention – a competition that needs to be resolved following anti, but not pro-retrocues (as also reported in van Ede et al, 2020). These findings replicate our earlier work and form the basis for the EEG results that were not available in our original study and that thus form the key advances of the current work.

### EEG

#### Cue-evoked activity: informative retrocues modulate the cue-evoked sensory response

In van Ede et al. (2020) and in this study (Fig. 2B), the eye tracking data suggest that informative retrocues prompt the internal selection of the to-be recalled memory item. Based on this, one might expect that the retrocue receives differential processing when it is informative (in the pro- and anti-retrocue conditions relative to the null-cue condition). To examine this possibility, we calculated the cue-locked event-related potentials (ERP) of two bilateral posterior electrodes (i.e., P7/8) and focused on components P1 and N1 (Eason et al., 1969; Mangun and Hillyard, 1991). As depicted in Figure 3, we observed significant effects of retrocue conditions in both the P1 and N1 components (F(2, 48) = 4.07, p = 0.023; F(2, 48) = 8.48, p < 0.001, respectively). The follow-up tests showed that the ERP for pro- and anti-cues became more negative than for the null-cue soon after cue onset, a difference that persisted during the initial positive-going deflection at around 70 to 130 msec (p = .030 and p = 0.013, respectively) and the first negative-going deflection that peaked around 130 to 200 msec (p = .002 and p = 0.011, respectively; Fig. 3).

**Figure 3.**
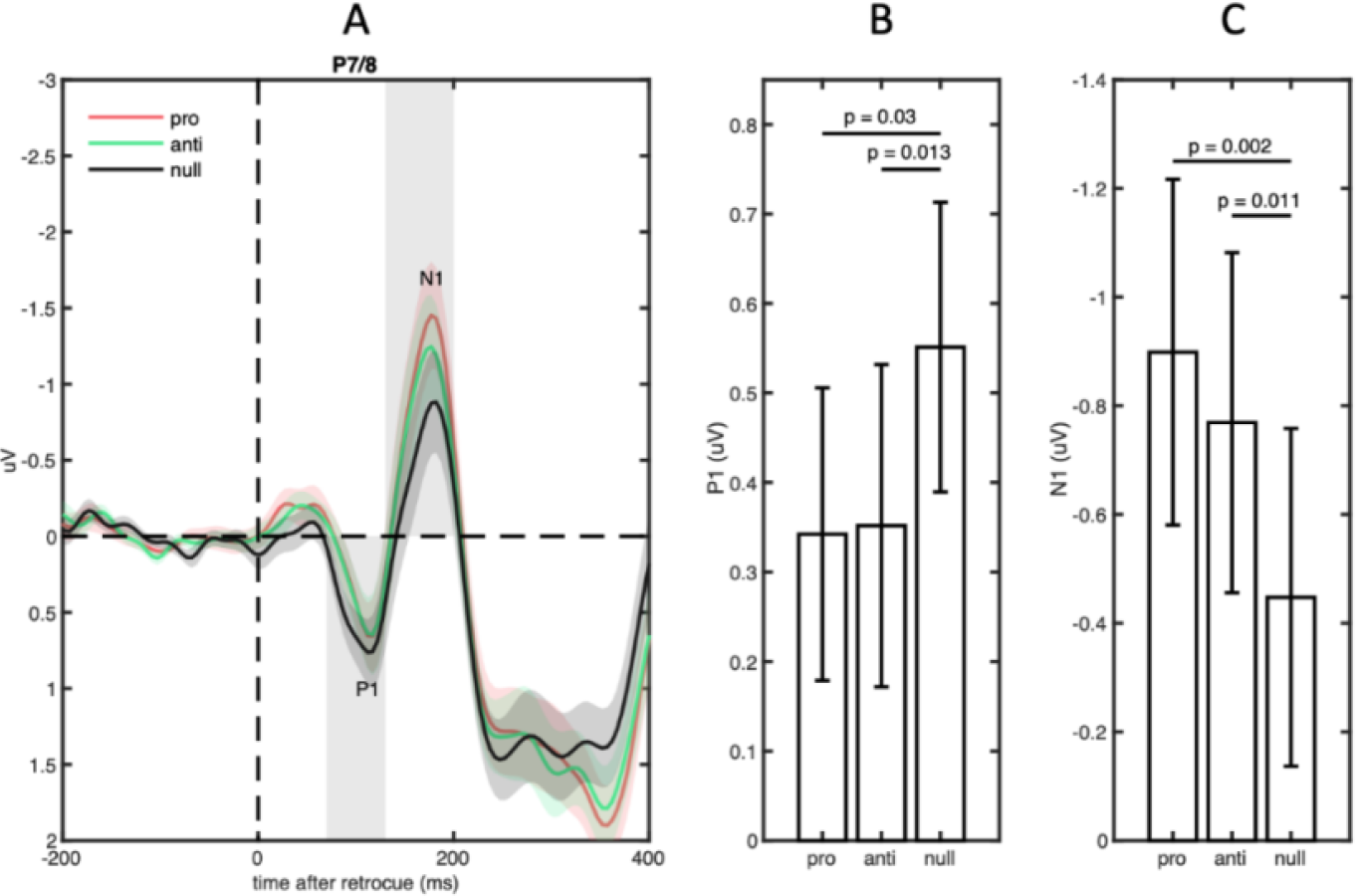
Cue-locked ERPs distinguish informative from uninformative retrocues. A. ERPs recorded from electrodes P7/8 indicate differential processing of informative retrocues. Shading represents ± 1 SEM. Vertical gray areas indicate time windows of interest to compare ERP components across conditions. B and C show mean ERP values of P1 and N1, respectively. Note how for the P1 we plotted positive upwards, while for the N1 we plotted negative upwards, such that a larger P1 or N1 response is associated with higher bars in both graphs. Error bars denote ± 1 SEM across observers.

#### Contralateral sensorimotor beta: informative retrocues induce action preparation and this is delayed following anti-retrocues

To investigate activity associated with action preparation, we assessed sensorimotor beta activity at electrode C3 (Fig. 4), which would emphasize activity related to the upcoming response hand (which was always the right hand in the current experiment). Compared to the null-cue condition, post-cue power in a frequency band spanning from approximately 9-15 Hz at C3, was decreased following pro retrocues beginning approximately 100 ms after cue onset (cluster P < 0.001, Fig. 4A, i), and on anti-cue trials beginning approximately 1000 ms after retrocue onset (cluster P = 0.007, Fig. 4B, i). In both conditions, this relative power attenuation persisted for the remainder of the second delay, and eventually incorporated a frequency band spanning approximately 8-30 Hz. The difference in the timing of these effects was confirmed by a significant difference between pro- and anti-cue trials during an epoch spanning from approximately 300-1300 ms after probe onset (cluster P = 0.014; Fig. 4C, i), and is consistent with the delay in the gaze bias following anti-compared to pro-retrocues. These action-preparation signals were most prominent over electrode C3 (Fig. 4, ii) consistent with an anticipated orientation-recall report with the right hand – a manual report that could be prepared for, at an abstract level, as soon as the appropriate item had been selected from working memory.

**Figure 4.**
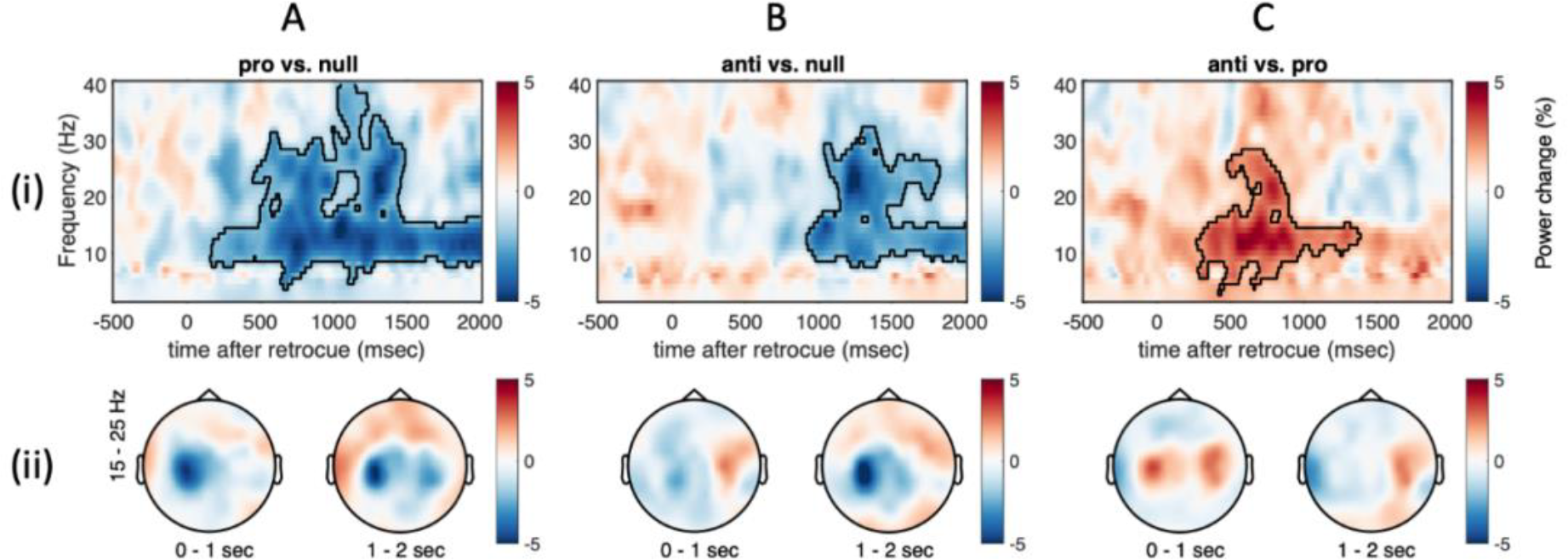
Informative retrocues initiate anticipatory action preparation that is delayed following anti-retrocues. (i) Comparison of time-frequency representations of oscillatory power, at electrode C3, for (A) pro-vs. null-retrocues, (B) anti-vs. null-retrocues, and (C) anti-vs. pro-retrocues. (ii) Topographies of the comparisons from row (i), restricted to the beta band (15-25 Hz) for two different time windows after retrocue onset. Colors indicate percentage differences between conditions; black cluster outlines indicate significant differences from a cluster-based permutation test.

#### Frontal midline theta: tracking demands on cognitive control over and above working-memory load

We now turn to our analysis focusing on frontal midline theta activity that was of key interest in the current work for reasons outlined in our introduction. Figure 5 shows spectral modulations at the frontal-midline electrode AFz following pro- and anti-retrocues, relative to following null-retrocues that here served as the condition to which to compare the EEG signal. Oscillatory power at AFz became significantly elevated across a range spanning from roughly 4-10 Hz following both pro- (cluster P = 0.004, Fig. 5A) and anti-retrocues (cluster P = 0.003, Fig. 5B), beginning approximately 800 ms after cue onset and persisting for the remainder of the second delay period.

**Figure 5.**
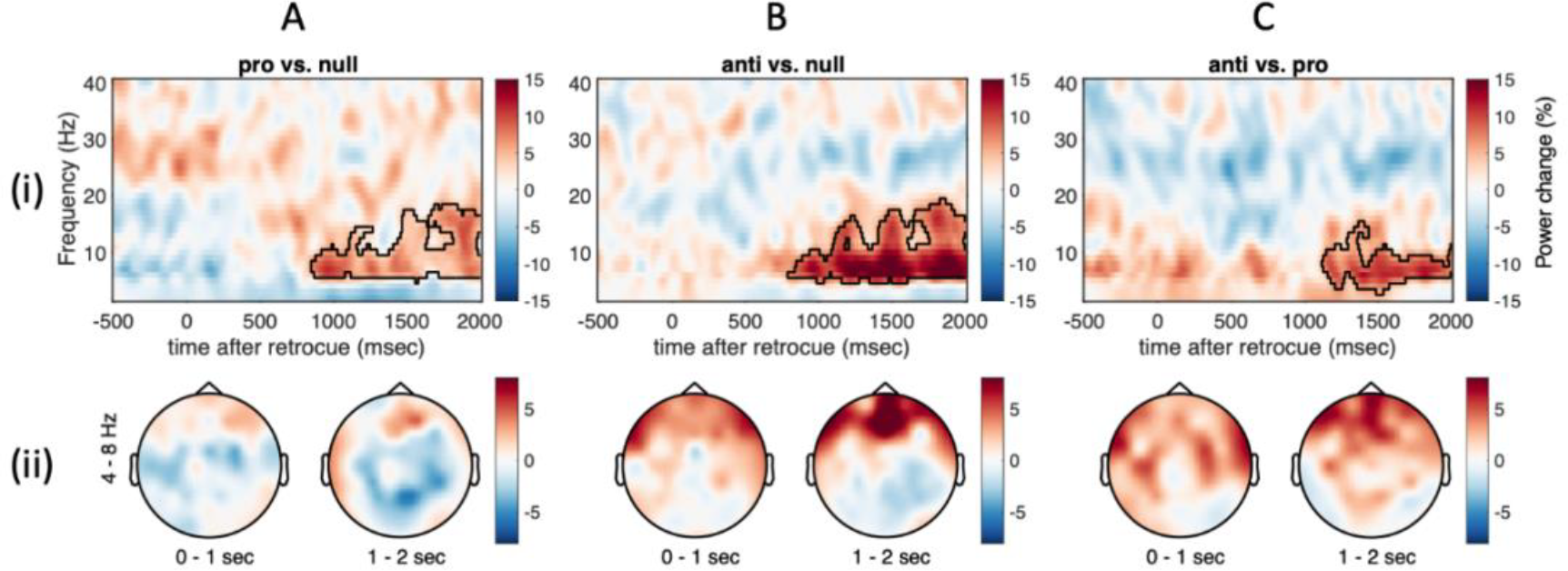
Frontal midline theta activity indexes demands on cognitive control. (i) Comparison of time-frequency representations of oscillatory power, at electrode AFz, for (A) pro-vs. null-retrocues, (B) anti-vs. null-retrocues, and (C) anti-vs. pro-retrocues, aligned to retrocue onset. (ii) Topographies of the comparisons from row (i), restricted to the theta band (4-8 Hz), for two different time windows relative to retrocue onset. Colors indicate percentage differences between conditions; black cluster outlines indicate significant differences from a cluster-based permutation test.

Note that, because the null-cue condition required the continued retention of two items while following the 100% informative retrocues load could be reduced to one item, these effects imply that the influence of selection demands on frontal midline theta may be greater than the influence of memory load.

In addition, when directly contrasting pro- and anti-retrocue conditions, we found larger theta modulations following anti-retrocues, consistent with larger demands on cognitive control following retrocues for which voluntary and involuntary factors compete for the focus of attention in working memory. This difference was most pronounced in the 4-10 Hz range that from roughly 1100 msec after cue onset (cluster P = 0.011, Fig. 5C).

#### Frontal midline theta predicts ensuing memory-guided behaviour

Finally, to explore how the above-described theta modulations might relate to behavior, we used a median split analysis to investigate how post-cue frontal midline theta-band power related to two aspects of behavior: response-initiation RT and recall precision. For response-initiation RT (Fig. 6), after performing a median split, we observed that theta power at AFz was numerically higher on fast relative to slow trials for all three conditions. For RT, this effect reached significance on both anti-retrocue trials (with the cluster beginning approximately 250 ms after retrocue onset; cluster p = 0.002, Fig. 6B), and on null-retrocue trials (with the cluster beginning approximately 450 ms prior to retrocue onset and extending beyond cue processing; cluster p < 0.001, Fig. 6C).

**Figure 6.**
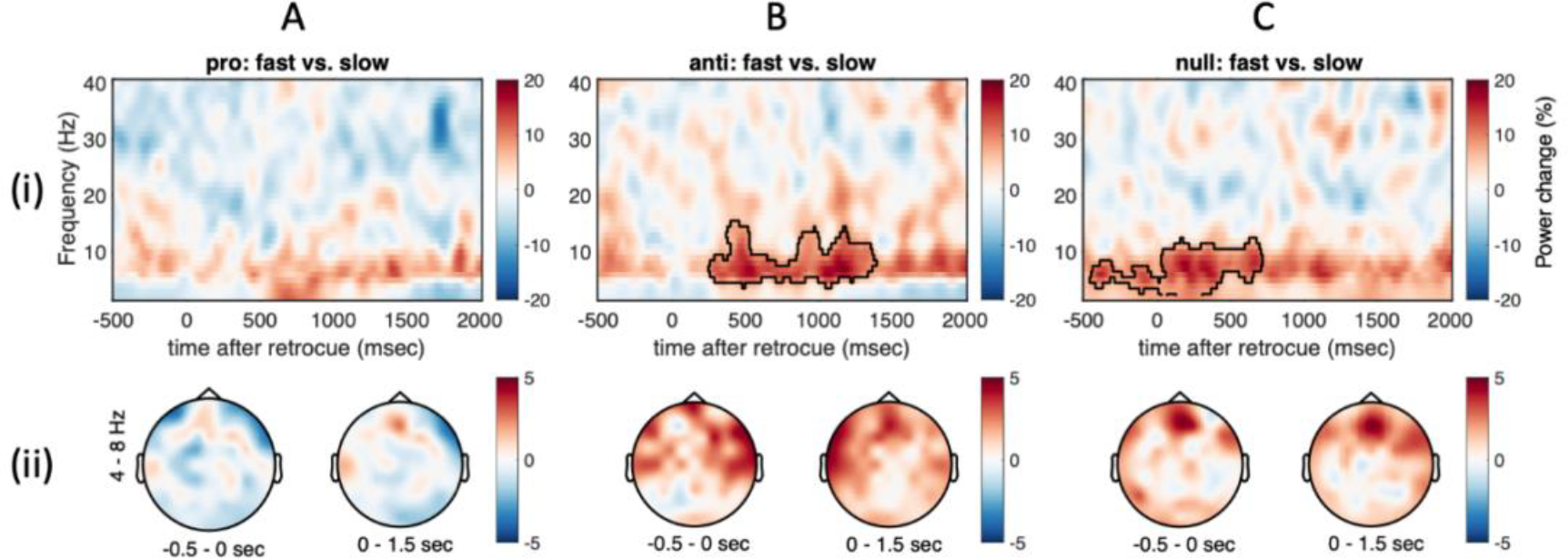
Frontal midline theta power predicts the response-initiation latency of ensuing working-memory-guided behaviour. (i) Comparison of time-frequency representations of oscillatory power, on fast versus slow trials (median split), at electrode AFz, for (A) pro-, (B) anti-, and (C) null-retrocue trials, aligned to retrocue onset. (ii) Topographies of the comparisons from row (i), restricted to the theta-band (4-8 Hz), for two different time windows relative to retrocue onset. Colors indicate percentage differences between conditions; black cluster outlines indicate significant differences from a cluster-based permutation test.

For recall precision, after performing a median split we also observed that delay-period oscillatory power at AFz was numerically higher on high-vs. low-precision trials for all three conditions, although these effects did not survive statistical significance testing (for transparency, we present these results in Appendix Figure A1).

## Discussion

Building on recent demonstration of joint voluntary and involuntary influences over the focus of attention inside visual working memory (van Ede et al., 2020), we set out to investigate the neural dynamics of such influences, and their competition. First, our results replicated the findings from van Ede et al. (2020) that performance improves with informative (pro- and anti-cue) relative to non-informative (null-cue) retrocues – confirming the successful manipulation of internal selective attention. We also replicated gaze biases associated with these internal attention shifts (c.f. van Ede et al., 2019; van Ede et al., 2020; Liu et al., 2022), and the delayed allocation of attention to appropriate memory content when voluntary and involuntary factors competed (van Ede et al., 2020). The concurrently recorded EEG extended these original findings in several ways. First, retrocue-locked posterior ERP components were shifted negatively and the N1 component boosted following informative (pro- and anti-retrocues) relative to uninformative null retrocues. Second, at the central electrode overlaying the left motor cortex, power centered on the mu-alpha and beta bands – indexing preparation for the ensuing manual report – decreased soon following pro-relative to null-retrocues, and markedly later following anti-relative to null-retrocues – paralleling the delayed attentional allocation seen in gaze, but here extending it to action planning alongside visual retention. Finally, post-retrocue frontal midline theta activity (FMT) was highest following anti-, then pro-, then null-retrocues, despite informative retrocues (but not null retrocues) allowing to reduce memory load from two to one. We consider each of these electrophysiological findings in turn.

Memory report was fastest and most accurate for pro-retrocue trials, then anti-retrocue trials, relative to null-retrocue trials. Although this report couldn’t begin until 1750 msec after cue onset, the cue-evoked ERPs suggest that the utilisation of informative retrocues may have begun with enhanced perceptual processing of the cue itself, presumably due to enhanced (voluntary) attentional gain modulation of the initial sensory response to the cues (c.f., Hillyard, Vogel, & Luck, 1998).

Subsequently, dynamics in the lower beta band at central electrodes indicated that quicker attentional allocation to the appropriate memory content afforded by pro-retrocues (as indexed by the quicker onset of gaze towardness) resulted in earlier engagement of anticipatory manual response preparation (c.f., Boettcher et al., 2021; Nasrawi & van Ede, 2022; Nasrawi et al., 2023; Schneider et al., 2017; van Ede, Chekroud, Stokes, et al., 2019). This shows that competition between voluntary and involuntary influences over the internal focus of attention results not only in delayed attentional allocation to appropriate memory content, but also in delayed initiation of preparation to act on this content (Rösner et al., 2022).

Turning to FMT, the findings are interesting from two perspectives: what they reveal about the control of competition for selection within visual working memory, and how they inform the interpretation of FMT dynamics as an index of working memory-related processing. Anti-retrocue trials were objectively more difficult than pro-retrocue trials. This was evidenced by RT and precision of recall, as well as by the delay in both the gaze bias (indexing attentional selection) and the central mu alpha/beta modulations (indexing action preparation). The dynamics of the gaze bias, in particular, suggest that anti-retrocues triggered an initial reflexive shift of attention toward the colour-matching item, an operation that would need to be overridden by rule-guided (i.e., voluntary) control of behavior (i.e., select the color-nonmatching item). FMT has long been associated with cognitive control (e.g., Sauseng et al., 2010; Cavanagh & Frank, 2014; Hsieh & Ranganath, 2014), and the fact that post-cue FMT was higher on anti-than pro-retrocue trials is consistent with a role in controlling the competition for selection (for a recent preprint that parallels this finding, see Ester & Nouri, 2022). Furthermore, the fact that post-cue FMT was also higher on pro-than null-retrocue trials suggest a more general role in the control of selection within working memory. Indeed, in two of the three conditions in this experiment we further observed that higher FMT following the cue predicted faster memory-guided behaviour, suggesting a functional role for these theta modulations.

How do the present results inform our understanding of working memory-related functions of FMT? Although to this point we have emphasized a role in indexing the need for control, it has also previously been reported that FMT power can track working-memory load (e.g., Gevins, 1997; Jensen & Tesche, 2002; Meltzer et al., 2008; Zakrzewska & Brzezicka, 2014; van Ede et al., 2013). Our design effectively pits the factor of control-of-selection versus that of load, because, in contrast to the pro- and anti-retrocue conditions, in which load could be reduced to a single item, the null-retrocue condition required retention of the two memory items throughout the post-cue delay. We observed higher FMT power following the retrocues that effectively triggered a reduction working-memory load. Consequently, our current results suggest that FMT cannot be interpreted as an index that is specific to working-memory load, per se. Instead, at a most general level, should be interpreted in the context of models of frontal midline systems (e.g., anterior cingulate cortex) involved in assessing the need for and regulation of the level of cognitive control (e.g., Shenhav et al., 2013; Araújo et al., 2002; Kahana et al., 1999; de Vries et al., 2019; Fiebelkorn & Kastner, 2019; Gevins, 1997; Gevins & Smith, 2000; Helfrich et al., 2018; Pennekamp et al., 1994; Sauseng et al., 2007; Vries et al., 2017).

One interesting observation with reference to our theta findings regards their timing. Although cluster statistics do not support inferences about the precise timing of events (Sassenhagen & Draschkow, 2019), it is nonetheless notable that the strongest most pronounced differences in FMT are in the period from 1000 to 2000 ms after retrocue onset, including the anti-vs pro-retrocue comparison (Figure 5). This is markedly later than the gaze and Central beta effects that indicated that the competition between voluntary and involuntary factors was resolved mostly within the first second after the retrocue. This opens the possibility that the pronounced FMT activity following anti-retrocues may not reflect the process of resolving competition per se, but rather the maintenance and/or consolidation of the post-cue state. It is important to keep in mind, however, that a different analysis of the same data (the correlation of FMT power with RT) revealed a significant functional role for FMT during the earlier portion of the post-cue delay in two conditions, and a trend in this direction for the third (Fig. 6). This is consistent with an important role for FMT in the flexible control of the selection and processing of the response-critical information in working memory.

In summary, we have used EEG to investigate the neural dynamics associated with the competition between voluntary and involuntary control over the focus of attention in visual working memory. When voluntary and involuntary factors compete for the internal visual focus of attention, prioritisation of the appropriate memory item is delayed. This is reflected both in gaze biases that track selection and in the neural dynamics of preparation for the appropriate upcoming manual action. We have further shown how this competition is paralleled – possibly followed – by frontal midline theta activity that influences memory-guided behavioural performance. Finally, our design uniquely enabled us to disentangle the processes of control versus retention in working memory, revealing how FMT tracks cognitive control demands over and above working memory load.

## Acknowledgements

This research was supported by an ERC Starting Grant from the European Research Council (MEMTICIPATION, 850636) and an NWO Vidi grant by the Dutch Research Council (grant number 14721) to F.v.E. and by National Institutes of Health (R01MH064498 and R01MH095984) to B.R.P.

## Appendix 1

**Figure A1.**
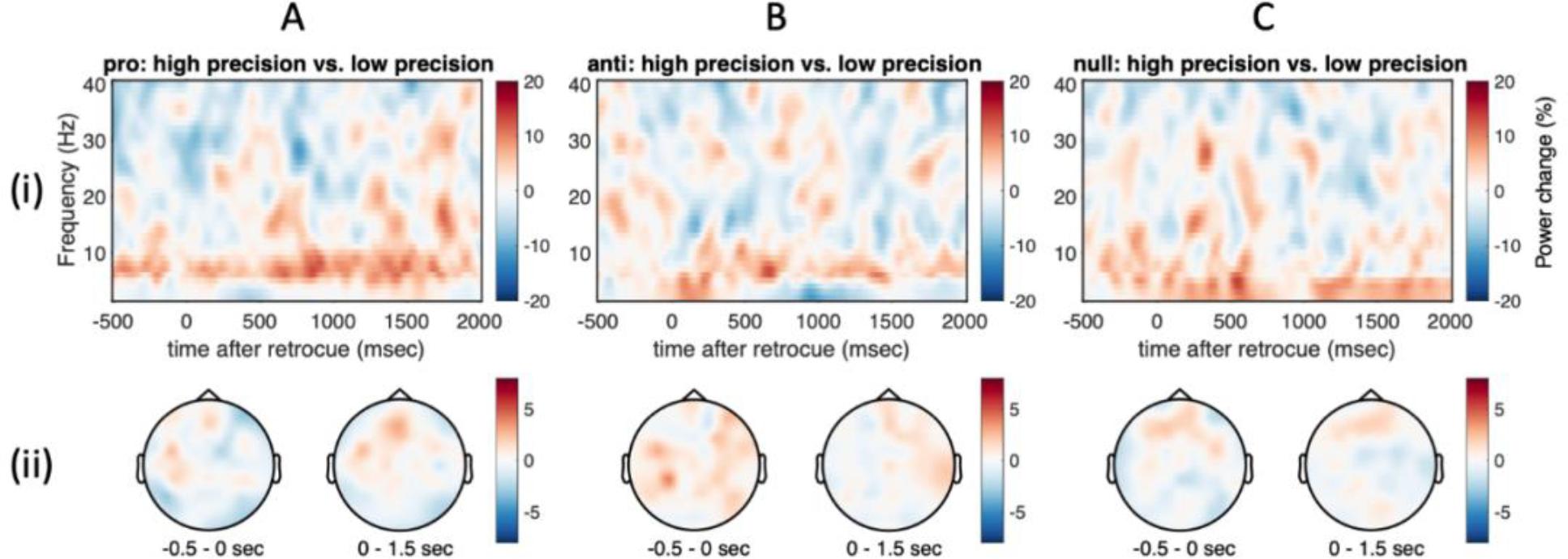
Comparison of neural activity between trials with high and low precision orientation reports separately for (A) pro, (B) anti, and (C) null retrocue trials. For each retrocue condition comparison: (i) difference in time-frequency response at AFz, aligned to retrocue onset. Colors indicate percentage differences between conditions. (ii) topographies of theta (4-8 Hz) percentage difference between high precision and low precision trials, for each retrocue condition comparison, for two different time windows after retrocue onset.

## Notes

### Competing Interest Statement

The authors have declared no competing interest.

